# Repeated Exposure Decreases Aesthetic Chills Likelihood but Increases Intensity

**DOI:** 10.1101/2024.03.01.582918

**Authors:** F. Schoeller, L. Christov-Moore, C. Lynch, A. Jain, T. Diot, N. Reggente

**Affiliations:** Institute for Advanced Consciousness Studies, Santa Monica, CA, USA; MIT Media Lab, Cambridge, MA, USA; Department of Psychiatry, 7th Sector, GHU Paris Psychiatrie et Neurosciences, Paris, France

**Keywords:** Aesthetic Chills, Habituation, Psychophysiology, Emotional Response, Reward Processing

## Abstract

Aesthetic chills are a peak emotional response to affectively charged stimuli such as music, films, or speech. This study investigates the impact of repeated exposure on the frequency and intensity of aesthetic chills. Through a longitudinal approach, we quantified changes in chill likelihood, intensity, and pleasure across multiple exposures, focusing on audio stimuli. Participants (n = 58) were randomly exposed to 6 chill-evoking stimuli pre-validated on the population of interest, in a counterbalanced order. Our findings revealed a significant decrease in the likelihood of experiencing chills with repeated exposure, suggesting habituation to chills itself or potential fatigue in response to aesthetic stimuli. The study also identified distinct demographic and psychophysiological response patterns across different participant groups, indicating variability in chill responses. These results provide insights into the dynamic nature of aesthetic experiences and their underlying neural mechanisms, with implications for understanding emotional and reward processing in psychophysiology.

## 1. Introduction

Aesthetic chills (hencefoth “chills”) are an intense pleasurable experience characterized by distinct physiological markers including piloerection and changes in skin conductance, heart rate, respiration, and body temperature (Schoeller et al., 2023; Panksepp, 1995; McCrae, 2007; Benedek & Kaernbach, 2011). Chills typically occur during peak emotional responses to stimuli like music, films, and speeches, engaging brain regions linked to reward/avoidance and dopamine release (Blood & Zatorre, 2001; Salimpoor et al., 2011). Recent evidence suggests chills may have therapeutic potential for disorders involving dopamine dysfunction like depression and anhedonia (Jain et al., 2023). However, existing chills research relies heavily on self-selected musical excerpts, limiting generalizability and introducing pre-selection biases (Patten, 2000). Crucially, the longitudinal dynamics of chills responses remain poorly characterized, hindering translation into personalized clinical applications. Peak emotional states like chills are susceptible to habituation, where repeated exposure attenuates the intensity of the response over time (Rankin et al., 2009). Quantifying such habituation effects constitutes a key step in developing interventions leveraging short-term alterations in consciousness for long-term clinical benefits.

In this first preliminary study, we assessed changes in chills experiences during repeated exposure to chills-evoking audiovisual stimuli (Schoeller et al., 2022, 2023). We leveraged a set of six pre-validated chill-evoking audiovisual stimuli, sourced from ecologically valid platforms (YouTube, Reddit) and empirically validated prior across 3500+ listeners (Schoeller et al., 2022). A focus was placed on music and speech given their engagement of reward and emotional circuits (Levitin & Tirovolas, 2009; Frühholz et al., 2016). To explore potential habituation effects to repeated chills exposure, we quantified longitudinal changes in chills likelihood and parameters (frequency, intensity, duration). We hypothesized that chills likelihood and intensity would decrease with repeated exposure due to habituation. Characterizing such dynamics constitutes a critical first step towards developing positive affect exposure interventions for disorders linked to blunted reward sensitivity.

## 2. Methods

### 2.1. Participants

58 participants from Southern California were recruited online using the Prolific platform. Most participants identified as White or Caucasian (63.79%, N=37) and had a Bachelor’s degree (50.0%, N=29). A common income level was “$75,000-$99,999” (22.41%, N=13). There was a fairly even mix of Male (50.0%, N=29) and Female (44.83%, N=26) participants, with a small number identifying as Non-Binary/Fluid (5.17%, N=3).

### 2.2. Stimuli

The stimuli utilized in this study were sourced from ChillsDB, a comprehensive database of chill-inducing stimuli that have been empirically validated across a sample of over 3000 participants from Southern California. Stimuli spanning various modalities (films, music, speech) were selected to ensure a wide-ranging exploration of chills, while also considering the varied chills ratio associated with each stimulus (table 1).

**Table 1:** The table provides details about the six different videos used in the study. For each video, a brief description, its length, and its chills ratio (CR) from past ChillsDB studies are included. The CR tells us how often chills were experienced by viewers in past studies. The descriptions give a quick idea about what each video is about and what viewers might experience.

| Stimulus | Description (including duration, chills ratio) |
| --- | --- |
| Hallelujah (6:26, CR=.7) | Choir! Choir! Choir! began as a weekly drop-in singing event, equal parts singing, comedy, and community-building. In 2015, 1500 singers came to Luminato Festival in Toronto for a very special event. Choir! leaders taught them backup parts to Leonard Cohen's Hallelujah, and Rufus Wainwright joined the crowd on stage to lead the vocals. The collective performance amplifies the power of the musical piece. |
| Great Dictator (4:11, CR=.58) | In the 1940 satire-drama The Great Dictator, Charlie Chaplin plays 2 doppelganger characters: a fascist dictator and a Jewish barber avoiding persecution. In this scene, the Barber is mistaken for the Dictator and must give a speech. Revered as one of the greatest monologues in film history, Chaplin makes a call for brotherhood and goodwill, encouraging soldiers to fight for liberty, and unite people in the name of democracy. What you are about to hear is the voice of Charlie Chaplin from the middle of World War II. For his entire career, he had been a silent actor. But in 1940, he decided to speak out. This is his message of hope. |
| Unbroken (5:57, CR=.56) | This motivational compilation, from a series by Youtube creator Mateusz M., combines quotes from Les Brown, Steve Jobs, Eric Thomas, and Louis Zamperini, with scenes from Her and Jobs to make a single motivational message. It is intended to encourage the listener to take full responsibility for one's life and follow one's path, even when it's unconventional; to pursue what one loves, and live a principled life in the face of hardship. |
| We think too much and feel too little (5:27, CR=.53) | This inspirational content, from Youtube creator Slyfer2812, combines scenes from a number of films (including Birdman, Chaplin, I am a legend, Into the Wild, Moonlight, The Greatest Showman, The Theory Of Everything, and Iron Man 3) to create a single motivational message. It is intended to encourage the audience to reflect on the importance of relationships and humans' profound connection to everything, inspiring not just to survive but to truly live. |
| Agnus Dei (7:22=.51) | The Flemish Radio Choir performs Samuel Barber's Agnes Dei, Opus 11, in Brussels. Agnus Dei ("Lamb of God") comes from John 1:29 and is a part of the Latin liturgical tradition. The words of the song are a request for peace and forgiveness. In 1967, Barber set the Latin words of the liturgical Agnus Dei, a part of the Mass, for mixed chorus with optional organ or piano accompaniment. |
| Silva (2:37, CR=.5) | Storyteller Jason Silva considers the impermanence of all and the dual nature of human consciousness, through a meditation on the a passage from Ernest Becker: "Man is literally split in two: he has an awareness of his own splendid uniqueness in that he sticks out of nature with a towering majesty, and yet he goes back into the ground a few feet in order blindly and dumbly to rot and disappear forever" -Ernest Becker |

### 2.3. Procedure

Participants were recruited via the Prolific platform, targeting individuals in Southern California. Upon recruitment, participants were redirected to a webpage specifically designed to host the experiment. The experiment began with the collection of demographic data, where participants responded to questions concerning their race, education level, household income, age, and sex, aligning with established demographic query standards. Following demographic data collection, participants were prompted to report their initial emotional and psychological state by responding to questions concerning their current valence, arousal, tension, and energy levels. Participants were then randomly exposed to one of the six chills stimuli. After engaging with the stimulus, participants were requested to answer questions pertaining to their chills experience, explicitly probing into the frequency, intensity, and duration of the chills experienced during the stimulus exposure. Additional questions concerning physiological responses (such as tears and goosebumps), as well as their overall enjoyment and liking of the video, were also posed to the participants. This cycle of stimulus exposure followed by post-stimulus questioning was repeated for all six stimuli. The order of stimuli presentation was randomized to control for potential order effects and to ensure the generalizability of the findings. After the final stimulus exposure and corresponding questions, participants were prompted to report their emotional and psychological state, capturing their valence, arousal, tension, and energy levels post-experiment. Concluding the experiment, participants responded to open-ended questions concerning their perspectives on the length and complexity of the questionnaire. Additionally, participants were provided with contact information through which to learn more about the chills phenomenon and the aims and hypotheses of the study. Participant feedback was predominantly positive, indicating good reception and engagement with the experiment.

### 2.4. Statistical analyses

Linear regression models were constructed to examine the relationships between presentation order and chills intensity, frequency, and duration. Models included presentation order as the predictor and the various chills parameters as outcomes. Standard regression diagnostics were performed, including analyses of residuals.

A multivariate logistic regression model was created to analyze the effects of presentation order, valence, and arousal on the likelihood of chills, with individual participant and specific stimuli coded as random effects. Model fit was assessed using Akaike’s Information Criterion (AIC). To categorize participants based on psychological response patterns, a hierarchical cluster analysis was conducted on trajectories of valence, arousal, tension, and energy ratings across stimulus presentations. Ward’s linkage method with squared Euclidean distances was used to generate clusters. Differences between clusters were analyzed via ANOVA and chi-squared tests. For all analyses, statistical significance was defined as p < .05. Estimates are reported alongside standard errors and test statistics. Effect sizes are indicated using R-squared values for regression analyses and odds ratios for the logistic regression. Plots visualizing key relationships supplement the analyses. The analyses were conducted using the R software (version 4.3.2).

### 2.5. Ethics

The experiment is in compliance with the Helsinki Declaration. Following review, the study protocol was granted anexemption status by Advarra IRB (Pro00068209). All participants gave their voluntary informed consent and we followed the Ethics Code of the American Psychological Association. All participants were informed about the purpose of the research, their right to decline to participate and to withdraw from the experiment, and the limits of confidentiality. We also provided them with a contact for any questions concerning the research and with the opportunity to ask any questions regarding the phenomenon under study (aesthetic chills) and receive appropriate answers.

## 3. Results

### 3.1. Chills likelihood decreases with presentation order

After creating several univariate models (not presented) for each of the four variables (valence, arousal, tension, and energy), we constructed a multivariate model using the Akaike Information Criterion (AIC) as a statistical criterion. This model incorporates the stimulus presentation order, valence, and arousal as significant factors. The likelihood of experiencing shivers decreased with the number of stimuli presented (Figure 1). Conversely, it increased when individuals exhibited higher levels of valence and arousal (p < 0.01). This result takes into account the difference between individuals and the various stimuli received by each.

**Figure 1.**
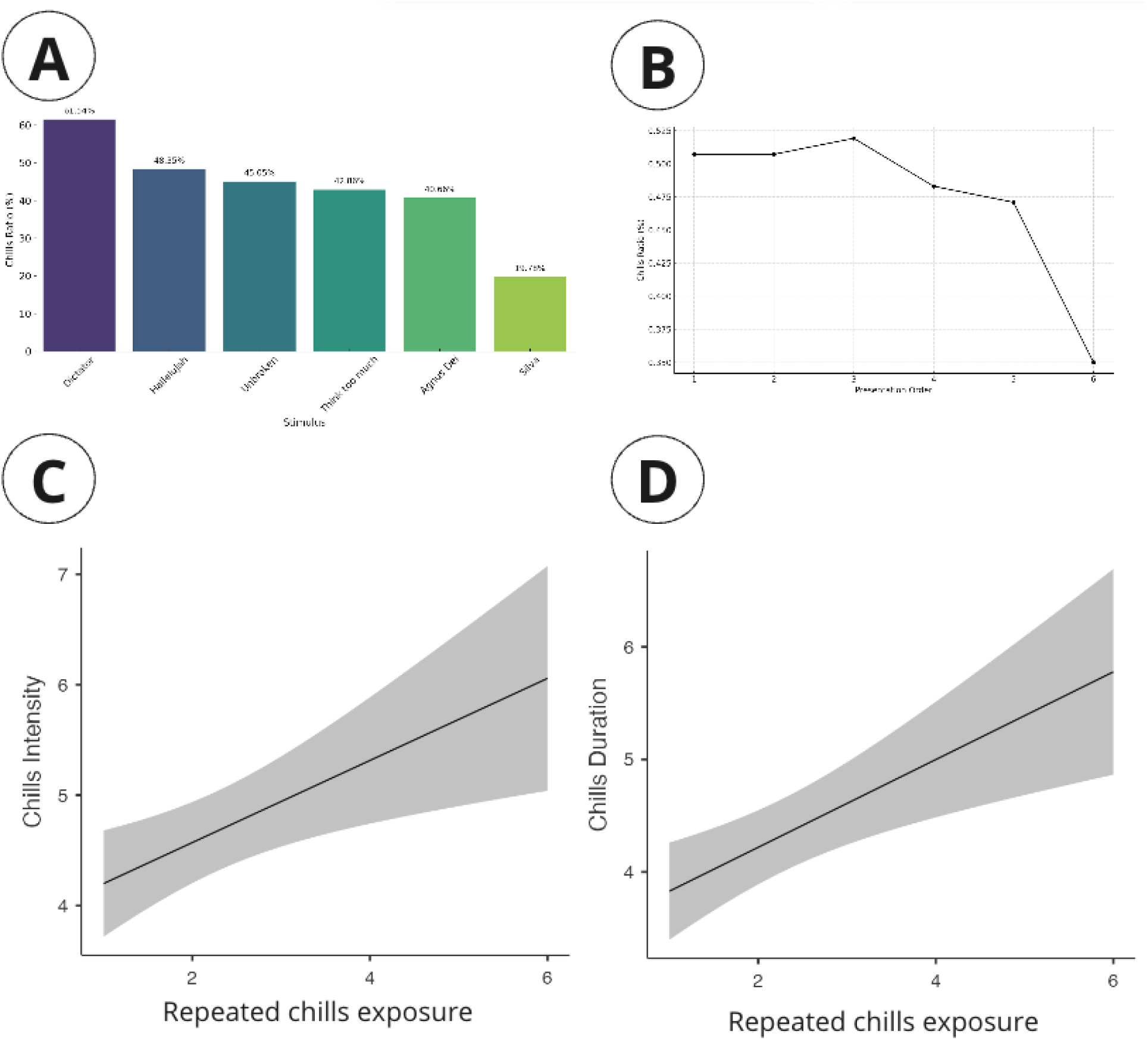
A: Bar chart showing the percentage of participants experiencing chills in response to various stimuli, ordered by descending frequency. B: Line graph indicating a decrease in the chills ratio over successive presentations of stimuli, with a marked drop between the fifth and sixth exposure. C: Plot illustrating the positive correlation between the intensity of chills and the number of repeated exposures to chills stimuli.D: Graph depicting an increase in the duration of chills with repeated exposure to the stimuli.

We conducted a linear regression analysis examining the relationship between presentation order and the intensity of chills among participants who experienced chills, a significant effect was found. The estimate for presentation order was 0.371 with a standard error of 0.129, resulting in a t-value of 2.89 and a p-value of 0.004. The intercept was also significant with an estimate of 3.829, a standard error of 0.343, and a t-value of 11.18 (p < .001). The model explained approximately 4.5% of the variance in chills intensity (R-squared = 0.0450), indicating a modest overall fit with a correlation coefficient of 0.212.

For chills frequency, the linear regression analysis indicated that presentation order did not significantly predict frequency (estimate = 0.0653, standard error = 0.0509, t = 1.28, p = 0.201). The intercept was significant (estimate = 1.8586, standard error = 0.1353, t = 13.73, p < .001). The model’s R-squared was 0.00918, suggesting that only about 0.918% of the variance in chills frequency was accounted for by the presentation order.

Regarding chills duration, the analysis showed a significant effect of presentation order (estimate = 0.390, standard error = 0.115, t = 3.37, p < .001). The intercept was significant as well (estimate = 3.440, standard error = 0.307, t = 11.19, p < .001). The model explained about 6.01% of the variance in chills duration (R-squared = 0.0601), with a correlation coefficient of 0.245 (Table 2).

**Table 2.** Multivariable mixed effect logistic regression for chills, with individual id and stimulus as a random effect. Individual intercept variance = 1.02 and stimuli intercept variance = 0.62 (Number of observations: 346. ID groups: 58; Stimulus groups: 6)

|  | **OR [95%CI] for chills** | **p-value** |
| --- | --- | --- |
| Intercept | 0.22 [0.06 ; 0.79] | **0.02** |
| Stimulus order | 0.77 [0.66 ; 0.9] | **< 0.01** |
| Valence | 1.26 [1.09 ; 1.46] | **< 0.01** |
| Arousal | 1.24 [1.09 ; 1.4] | **< 0.01** |

### 3.2. Three Distinct Groups in Chill Response

The analysis of trajectories of psychological parameters before and after each stimulus identified three distinct groups (Figure 2, Table 3). The first group (cluster A) is characterized by an average level of chills, with the lowest levels of valence and energy, and the highest level of tension. This group is primarily composed of women (p < 0.01) with a high level of education. Additionally, this group stands out for having the highest average duration of chills before any stimulation. The second group (cluster B) exhibits the lowest rate of chills, with low levels of tension and arousal and a high level of energy. This group consists mostly of men with a low level of education. They also had the shortest average duration of chills before stimulation. The last group (cluster C) displays the highest rate of chills during the experience and is characterized by very high levels of valence, arousal, and energy. This group comprises older white men (average age 45) with a relatively high level of education.

**Table 3:** Comparative Analysis of Chills Experience Across Three Groups: This table presents a detailed comparison of three groups (A, B, and C) in terms of their experiences with chills. It includes the number of participants in each group, frequency, intensity, and duration of chills, along with demographics such as gender, age, race, education level, and income. Statistical significance (p-value) is also provided for each category.

|  | A (n = 29) | B (n = 16) | C (n = 13) | p |
| --- | --- | --- | --- | --- |
| Number of chills |  |  |  | 0.02 |
| Mean (SD) | 2.5 (1.5) | 2.1 (1.5) | 3.8 (1.8) |  |
| Median [Q1, Q3] | 2.0 [2.0, 3.0] | 2.0 [1.5, 3.2] | 4.0 [3.0, 5.0] |  |
| **Gender** |  |  |  | **0.01** |
| Female | 18 (62.1%) | 4 (25.0%) | 4 (30.8%) |  |
| Male | 8 (27.6%) | 12 (75.0%) | 9 (69.2%) |  |
| Non-Binary/Fluid | 3 (10.3%) | 0 (0.0%) | 0 (0.0%) |  |
| **Age** | 33.8 (12.7) | 34.0 (11.1) | 44.5 (14.6) | **0.04** |
| **Races that you consider yourself to be** |  |  |  | 0.49 |
| Black or African American, Asian, Other | 5 (17.2%) | 6 (37.5%) | 2 (15.4%) |  |
| White + Other | 5 (17.2%) | 1 (6.2%) | 2 (15.4%) |  |
| White or Caucasian | 19 (65.5%) | 9 (56.2%) | 9 (69.2%) |  |
| **Highest level of education** |  |  |  | **0.02** |
| Associates or technical degree | 1 (3.4%) | 2 (12.5%) | 4 (30.8%) |  |
| Bachelor's degree, graduate or professional degree (MA, MS, MBA, PhD, JD, MD, DDS etc.) | 19 (65.5%) | 7 (43.8%) | 6 (46.2%) |  |
| High school diploma or GED | 7 (24.1%) | 1 (6.2%) | 2 (15.4%) |  |
| High school or some college, but no degree | 2 (6.9%) | 6 (37.5%) | 1 (7.7%) |  |
| **Total household income before taxes during the past 12 months** |  |  |  | 0.96 |
| Less than \$25,000 | 5 (17.9%) | 3 (18.8%) | 2 (15.4%) | |
| \$25,000-\$49,999 | 4 (14.3%) | 4 (25.0%) | 2 (15.4%) | |
| \$50,000-\$74,999 | 8 (28.6%) | 1 (6.2%) | 3 (23.1%) | |
| \$75,000-\$99,999 | 6 (21.4%) | 4 (25.0%) | 3 (23.1%) | |
| \$100,000-\$149,999 | 3 (10.7%) | 2 (12.5%) | 2 (15.4%) | |
| \$150,000 or more | 2 (7.1%) | 2 (12.5%) | 1 (7.7%) | |
| **Chills frequency** |  |  |  | 0.20 |
| More than Once a week | 14 (48.3%) | 3 (18.8%) | 3 (23.1%) |  |
| Never or less than Once a year | 4 (13.8%) | 3 (18.8%) | 4 (30.8%) |  |
| Once a month | 11 (37.9%) | 10 (62.5%) | 6 (46.2%) |  |
| **Chills intensity** | 5.8 (1.5) | 5.0 (1.9) | 5.3 (2.5) | 0.40 |
| **Chills duration** | 3.7 (1.3) | 2.6 (1.0) | 3.4 (1.6) | **0.03** |
| **Chills like** | 5.4 (2.5) | 5.6 (2.6) | 5.1 (2.8) | 0.88 |

**Figure 2.**
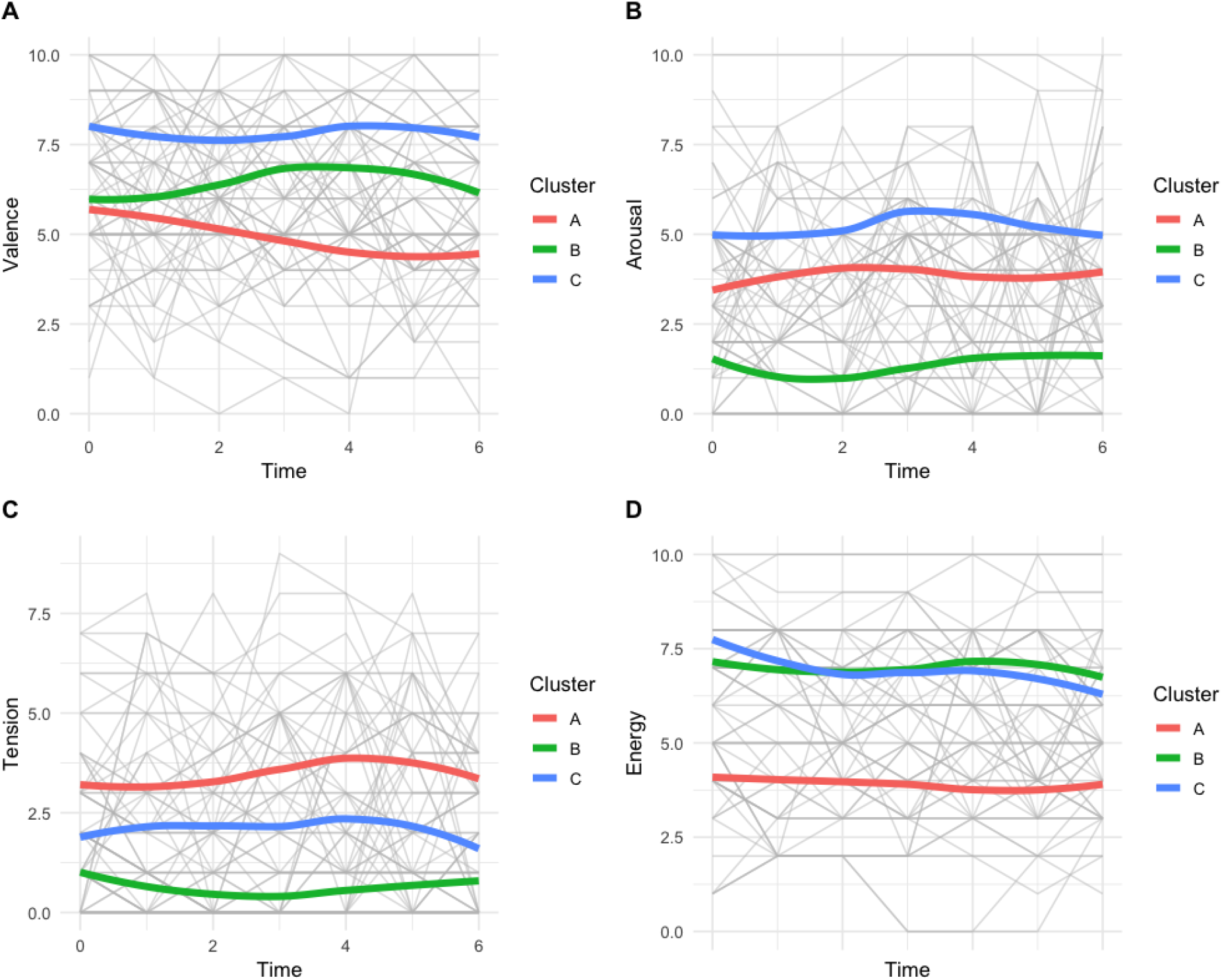
Evolution of the four pre-stimuli psychological parameters by cluster (A: Valence, B: Arousal, C: Tension, D:Energy). Each color represents one cluster, and the grey line represents individual evolution.

## 4. Discussion

We found a decline in chills likelihood with repeated stimulus exposure. Specifically, a decreasing trend in chills likelihood across stimulus presentations, with a notable drop from the fifth to the sixth stimulus. This suggests potential habituation of the neurophysiological mechanisms supporting chills, consistent with existing evidence on the effects of repeated exposure on other such reward responses (Dreyer et al., 2010). By this interpretation, the observed decline likely stems from dynamic dopaminergic modulation over repeated exposures, gradually reducing excitability of reward pathways. Potential neural habituation mechanisms include dopamine receptor internalization, depletion of readily releasable dopamine vesicular stores, and adaptations in postsynaptic signaling cascades (Borland & Michael, 2004). Elucidating the precise habituation pattern and neural underpinnings can shed light on aesthetic emotion dynamics and reward learning.

Beyond habituation effects, our results also underscore the relationship between chills and pre-stimulus psychological states. We found that higher valence and arousal levels correlate with increased chills likelihood. This aligns with prior evidence linking chills to physiological arousal and positive emotions (Benedek & Kaernbach, 2011; Panksepp, 1995). Our analysis of psychological trajectories revealed groups exhibiting positive arousal changes post-listening also had higher chill frequencies. This further supports chills’ role as a marker of strong emotive responses to aesthetic stimuli. Notably, while the overall sample showed habituation effects, a subset of participants experienced consistent chills with heightened intensity across repetitive exposures. This points to meaningful individual differences in reward pathway excitability and aesthetically-driven emotional reactivity. Integrating neural and phenotypic data could elucidate if such sensitivity stems from endogenous factors like genotype or sociocultural exposures.

At a cellular level, potential habituation of the chills response may be underpinned by dopaminergic adaptations that modulate reward sensitivity over repeated exposures. Such mechanisms could include dopamine receptor internalization, depletion of readily releasable vesicular dopamine stores, or changes in postsynaptic signaling cascades (Borland & Michael, 2004; Dreyer et al., 2010). Elucidating the rate of decay of the chills response and its neural correlates can provide key insights into these habituation processes and their implications for reward learning and aesthetic emotion.

## 5. Conclusion

This study provides initial evidence of declines in chills likelihood with repeated aesthetic stimulus presentation. This likely habituation effect may stem from dynamic modulation of dopaminergic reward pathways. Elucidating these neural habituation patterns could inform the design of chills-evoking therapeutic interventions for disorders like anhedonia and depression. Further integrating behavioral, neural, and genomic data could unveil the roots of such aesthetic emotion sensitivity. Overall, quantifying chills dynamics helps characterize an intriguing facet of human aesthetic appreciation and its ties to wellbeing.

## 6. Declaration

### Author contributions

Conceptualization: FS, NR, AJ

Methodology: FS, NR, AJ

Investigation: FS, NR, AJ

Visualization: FS, TD

Analysis: FS, TD

Writing: all

### Data and materials availability

All data are available in the main text or the supplementary materials. The ChillsDB 2.0 dataset is released under a Creative Commons Attribution 4.0 International (CC BY 4.0) license on FigShare (Schoeller et al., 2023). This allows others to freely share and adapt the dataset as long as appropriate credit is given to the original creators by citing the published paper (Schoeller et al., 2023). The dataset is divided into two .csv files available under a CC BY 4.0 license on the associated FigShare (ChillsDB2). For a comprehensive understanding of each column, researchers are advised to refer to the Header Explanation File. All code supporting these analytical efforts is included in the following repository. Note: Requires the LibSVM toolbox.

https://github.com/Institute-for-Advanced-Consciousness/E4-F01

All the data is accessible at the following FigShare:

Schoeller, Felix; Christov-Moore, Leonardo; Lynch, Caitlin; Jain, Abhinandan; Diot, Thomas; Reggente, Nicco (2024). Repeated Exposure Decreases Aesthetic Chills Likelihood but Increases Intensity. figshare. Dataset. https://doi.org/10.6084/m9.figshare.25000364.v1

## References

Benedek, M., & Kaernbach, C. (2011). Physiological correlates and emotional specificity of human piloerection. Biological Psychology, 86(3), 320–329.

Blood, A. J., & Zatorre, R. J. (2001). Intensely pleasurable responses to music correlate with activity in brain regions implicated in reward and emotion. Proceedings of the National Academy of Sciences, 98(20), 11818–11823.

Borland, L. M., & Michael, A. C. (2004). Voltammetric study of the control of striatal dopamine release by glutamate. Journal of neurochemistry, 91(1), 220–229.

Dreyer, J. K., Herrik, K. F., Berg, R. W., & Hounsgaard, J. D. (2010). Influence of phasic and tonic dopamine release on receptor activation. Journal of neuroscience, 30(42), 14273–14283.

Frühholz, S., Trost, W., & Kotz, S. A. (2016). The sound of emotions-Towards a unifying neural network perspective of affective sound processing. Neuroscience & Biobehavioral Reviews, 68, 96–110.

Jain, A., Horowitz, A. H., Schoeller, F., Leigh, S. W., Maes, P., & Sra, M. (2023). Reward Learning and Hedonic Experience in Major Depressive Disorder: Assessing the Potential of Aesthetic Chills as a Novel, Nonpharmacological Intervention. Journal of affective disorders, 302, 197–207.

Levitin, D. J., & Tirovolas, A. K. (2009). Current advances in the cognitive neuroscience of music. Annals of the New York Academy of Sciences, 1156(1), 211–231.

Mallik, A., Chanda, M. & Levitin, D. Anhedonia to music and mu-opioids: Evidence from the administration of naltrexone. Sci Rep 7, 41952 (2017). 10.1038/srep41952

McCrae, R. R. (2007). Aesthetic chills as a universal marker of openness to experience. Motivation and Emotion, 31(1), 5–11.

Patten S. B. (2000). Selection bias in studies of major depression using clinical subjects. Journal of clinical epidemiology, 53(4), 351–357. 10.1016/s0895-4356(99)00215-2

Panksepp, J. (1995). The emotional sources of” chills” induced by music. Music perception: An interdisciplinary journal, 13(2), 171–207.

Rankin CH, Abrams T, Barry RJ, Bhatnagar S, Clayton DF, Colombo J, Coppola G, Geyer MA, Glanzman DL, Marsland S, McSweeney FK, Wilson DA, Wu CF, Thompson RF. Habituation revisited: an updated and revised description of the behavioral characteristics of habituation. Neurobiol Learn Mem. 2009 Sep;92(2):135–8. doi: 10.1016/j.nlm.2008.09.012. Epub 2008 Nov 6. PMID: 18854219; PMCID: PMC2754195.

Schoeller, F. (2015). Knowledge, curiosity, and aesthetic chills. Frontiers in psychology, 6, 1546.

Schoeller, F., Christov-Moore, L., Lynch, C., Jain. A., & Reggente, N. (2023). ChillsDB 2.0: Individual Differences in Aesthetic Chills Among 2,900+ Southern California Participants. Open Science Framework. 10.31234/osf.io/s9qvk

Schoeller, F., Jain, A., Horowitz, A. H., Yan, G., Hu, X., Maes, P., & Salomon, D. Roy. (2022). ChillsDB, A gold standard for aesthetic chills stimuli. 10.31234/osf.io/9wrmq

